# Community lifespan, niche expansion and the evolution of interspecific cooperation

**DOI:** 10.1101/2020.01.02.893321

**Authors:** António M. M. Rodrigues, Sylvie Estrela, Sam P. Brown

## Abstract

Natural selection favours individuals who maximise their own reproductive success and that of their close relatives. From this perspective, cooperation that benefits individuals of a different species represents an evolutionary conundrum. The theory of mutualism seeks to resolve this puzzle, and it posits that there must be downstream benefits to cooperators that offset any costs inherently associated with interspecific cooperation. Thus, individuals should only further the survival and fecundity of their interspecific partners if this generates additional return benefits, such as food, shelter or protection. A major challenge for the evolution of mutualism is when the ecological niches of partner species overlap, as this creates a tension between the benefits of exchanging services and the costs of competing for shared resources. Here we study the extent to which niche expansion, in which cooperation augments the common pool of resources, can resolve this problem. We find that niche expansion facilitates the evolution of mutualism, especially when populations are at high densities. Further, we show that niche expansion can promote the evolution of reproductive restraint, in which a focal species adaptively sacrifices its own growth rate to increase the density of partner species. We interpret these results in the context of microbial community interactions, which are often characterized by yield-enhancing exchanges of nutrients, termed ‘cross-feeding’. Our findings suggest that yield-enhancing mutualisms are more prevalent in stable habitats with a constant supply of resources, where populations typically live at high densities, but such mutualisms are particularly vulnerable to the emergence of cheats. In general, our findings highlight the need to integrate both temporal and spatial dynamics in the analysis of mutualisms.

## Introduction

Multi-species communities are more than just an aggregate of individuals struggling for survival in the face of antagonistic interactions. While competitive and exploitative interactions are most common (e.g. Foster and Bell 2012), spectacular examples of interspecific mutualism enliven our natural world (Leigh 2010). On the microbial scale, examples of costly investments in the growth of a partner species have recently been described among microbes in the gut (Rakoff-Nahoum et al. 2016) and among partner species engineered (Shou et al. 2007, Wintermute and Silver 2010, Mee et al. 2014) and evolved in the lab (Harcombe 2010, Pande et al. 2014, Harcombe et al. 2018). These examples pose the question of whether costly microbial mutualisms are more widespread and awaiting discovery. To approach this question, we ask what ecological conditions are most likely to support these mutualistic investments?

Microbial cells release compounds into their environment that are then used by other members of the community for survival and growth (McNally and Brown 2015, Estrela et al. 2019). In some cases (termed cross-feeding or syntrophy), these exo-molecules are a waste product of an individual’s metabolism that is advantageously co-opted by a partner species (Hillesland and Stahl 2010, Estrela et al. 2012, LaSarre et al. 2017, Goldford et al. 2018), and as such, it comes at no cost to the producer. More challenging for evolutionary theory is when the exchanged compounds are not simple metabolic waste products but specifically designed costly secretions, such as the secretion of nutrient-scavenging molecules (e.g. siderophores; D’Onofrio et al. 2010) and digestive enzymes (e.g. glycoside hydrolase; Rakoff-Nahoum et al. 2016), or the production of detoxifying enzymes (e.g. beta-lactamases; Yurtsev et al. 2016). In the majority of these cases, the most parsimonious explanation is that the secreted factor serves to aid a conspecific (or even the focal producer cell), posing little dilemma if the anticipated recipient likely carries a copy of the gene coding for the trait. However, there are exceptions where the costly secreted factor provides benefits that are targeted to another strain or species (Harcombe 2010, Pande et al. 2014, Rakoff-Nahoum et al. 2016, Harcombe et al. 2018).

The interspecific exchange of costly secretions seems to be at odds with the Darwinian theory of adaptation by natural selection, which posits that selfishness should be rife in the natural world (Hamilton 1964a, b, Price 1970). The resources that an individual could use for its own selfish growth and survival are instead redirected to the production of exo-molecules that do not benefit their producer directly. Therefore, the evolution of these costly traits requires special conditions (Trivers 1971, Sachs et al. 2004, Foster and Wenseleers 2006, Leigh 2010). Mutualism theory provides the primary framework for understanding the adaptive nature of interspecific costly cooperation, where the general hypothesis is that the cost of interspecific cooperation must lead to a return benefit to the cooperator that offsets a cooperator’s initial costly investment (Trivers 1971).

Mutualisms, however, can become unstable when cheats can reap the benefits of interspecific cooperation without paying its costs. The stabilisation of mutualisms, therefore, requires the existence of specific mechanisms such that the return benefits reward cooperators rather than free riders. Three general stabilisation mechanisms have been proposed. First, partner fidelity occurs whenever individuals form close-knit lifetime associations, and therefore any products of cooperation are used up by partners, and not by selfish outsiders (Bull and Rice 1991, Sachs et al. 2004, Momeni et al. 2013). Second, partner choice enables cooperators to engage preferentially with other cooperators and to ostracise free riders (Bull and Rice 1991, Simms and Taylor 2002, Sachs et al. 2004, Regus et al. 2014). Finally, sanctions impose additional costs on cheats that offset any of the benefits associated with cheating (West et al. 2002, Kiers et al. 2003, Jandér and Herre 2010, Jandér et al. 2012, Regus et al. 2014).

Studies of microbial communities, however, have drawn attention to additional demographic challenges in the understanding of the adaptive exchange of molecules that have not been the focus of the general theory of mutualism (e.g. Foster and Bell 2012, Mitri and Foster 2013). Microbial communities have complex life histories where exchanges of services or resources occur while the community grows and depletes local resources, while it disperses and colonises new patches, or while the community experiences environmental disturbances. This rich ecological and demographic substrate provides the backdrop against which cooperative exchanges unfold, mediating the fitness costs and benefits associated with cooperation, and ultimately influencing the evolution of mutualism within microbial communities. For instance, theory suggests that interspecific cooperation is more likely to evolve at intermediate densities when individuals can reap the benefit of interspecific cooperation while avoiding the costs of interspecific competition (Bull and Harcombe 2009). This work, however, assumes a rate mutualism in which interspecific partners exchange resources that only influence their growth rates. Natural populations, by contrast, are often characterised by niche expansion in which individuals profit from an enlarged resource pool created by their interspecific partners (Samuel and Gordon 2006, Kolenbrander 2011, Ramsey et al. 2011, Goldford et al. 2018).

With a particular emphasis on cross-feeding interactions, here we study how different aspects of ecology, demography and genetic structuring mediate the evolution of mutualistic interactions. First, we focus on a rate mutualism and explore how the initial density of individuals, the lifespan of communities, local competition and relatedness within species interact with each other to mediate the evolution of mutualisms. Second, we extend our model to consider the possibility of niche expansion where interspecific partners expand the common pool of resources locally available. Specifically, we include cases in which niche expansion increases the local carrying capacity, and we determine how this, in turn, mediates the evolution of mutualisms.

## Model and Methodology

### Eco-evolutionary dynamics

We assume a metapopulation where each patch is colonised by *d*_0_ individuals of the focal species *A*, and by an equal number *d*_0_ of individuals of the partner species *B*. Interactions between the two species influence their growth trajectories, which we model using standard logistic equations. The intrinsic growth rate of species *A* is *ϕ*_A_(*z*), where *z* is the amount of help a focal individual of species *A* provides to individuals of species *B*. The intrinsic growth rate of the partner species *B* is *ϕ*_B_. Each individual of species *A* improves the growth rate of species *B* by an amount *β*(*z*). Similarly, each individual of the partner species *B* improves the growth rate of the focal species *A* by an amount *α*. There is density-dependent regulation in each patch, whose carrying capacity is *K.* The population of our focal species is composed of a resident strain that expresses a phenotype *z*, and of a mutant strain that expresses the phenotype *x* = *z* + *δ*, where *δ* is vanishingly small. Thus, the mutant strain deviates only slightly from the resident strain in their level of investment in helping behaviours. The growth trajectories of each species, including the resident and mutant strain of species *A*, is then given by

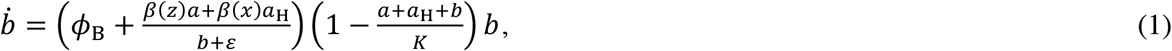

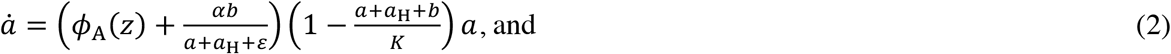

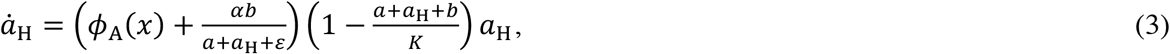

where: *b* is the density of the partner species *B*; *a* is the density of the resident strain of species *A*; *a*_H_ is the density of the mutant strain of species *A*; and < is a constant that mediates the growth rates of each species. After the initial colonisation of a focal patch, each species grows during a period of time *τ*, at which point there is competition for a new round of patch colonisation events. Within each species, we assume that a proportion *s* of the competition occurs locally, while a proportion 1 – *s* of the competition occurs globally, where *s* is the scale of competition (Frank 1998, Rodrigues and Gardner 2013). Individuals within each species compete for the *d*_0_ breeding sites available to each species in each patch, and those who fail to obtain a breeding site die. After this, the life-cycle of the community resumes (see Figure 1 for a schematic representation of the life-cycle).

**Figure 1.**
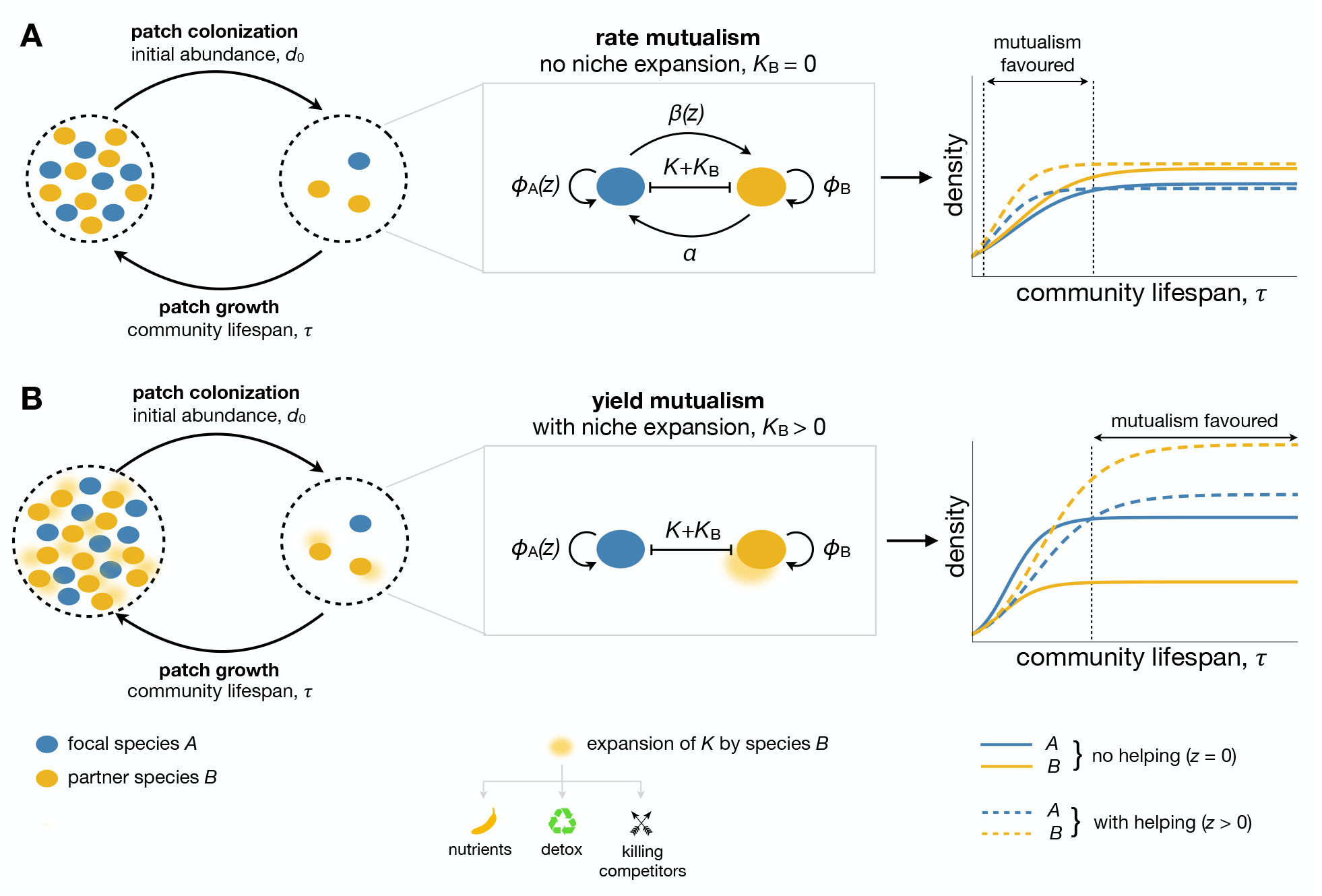
Model of the community lifecycle and interactions between species. We assume a metapopulation model where each patch is colonized by *d*_0_ individuals of the focal species *A* and of partner species *B*. Within each patch, species interact and grow until they disperse to a new empty patch. The carrying capacity of the patch is *K*. Two mutualistic interaction scenarios are illustrated: (**A**) a case of rate mutualism where the focal species *A* pays a cost on its own intrinsic growth rate (*ϕ*_A_(*z*)) to enhance the growth rate of its partner species *B* by *β*(*z*), and in return, gets a growth rate benefit *α* from *B*. This is defined as a rate mutualism because interspecific cooperation causes the two species to grow faster at intermediate stages of community development (here *K*_B_ = 0); (**B**) a case of yield mutualism in the form of reproductive restraint in the focal species *A* and niche expansion by *B*. Here the focal species *A* grows more slowly, which promotes the growth of the partner species *B*, and consequently increases the carrying capacity of the local patch by *K*_B_ (here *α* = *β*_0_ = 0). This is defined as a yield mutualism because interspecific cooperation causes both species to reach a higher equilibrium density.

### Fitness and stable strategies

We focus on helping that has a cost to the growth rate of individuals of species *A* and a benefit to the growth rate of the partner species *B*. We assume that the growth rate *ϕ*_A_(*z*) of species *A* is a decreasing function of the investment in helping *z*. Thus, d*ϕ*_A_(*z*)/d*z* < 0. In addition, we assume that the growth rate *β*(*z*) of species *B* is an increasing function of the investment in helping *z*. Thus, d*β*(*z*)/d*z* > 0 (see Figure S1 for a visual representation of the cost and benefit functions). Our aim is to understand how much helping *z* should individuals of the focal species *A* allocate to their partner species *B*. In other words, we want to determine the evolutionarily stable helping strategy *z*^*^, which is the strategy that cannot be beaten by an alternative strategy *z*^*^ ± *δ* (Otto and Day 2007).

To determine the stable strategy *z*^*^ of a focal individual of our focal species *A*, we need to define her reproductive success, which we denote by *f*(*x*,*z*), where the first argument *x* denotes the helping strategy of the focal strain and the second argument *z* denotes the helping strategy of the competing strain within the focal patch. We define the reproductive success of a focal individual as the number of descendants she produces after one growth round after she first colonises a patch, which is given by the final density of individuals in the local patch divided by the initial density of individuals in the same focal patch. Thus, the reproductive success of a focal individual that expresses the resident strategy is *f*(*z*,*x*) = *a*(*τ*)/(*d*_0_(1-*p*)), whilst the reproductive success of a focal individual that expresses the alternative strategy is *f*(*x*,*z*) = *a*_H_(*τ*)/(*d*_0_*p*), where *p* is the fraction of species *A* colonisers that are mutant individuals. A fraction *s* of the competition is local, in which case the focal individual competes with both clones of herself and wild-type individuals. The other fraction 1 – *s* of the competition is global, in which case the focal individual competes with wild-type individuals only. Thus, the fitness of a focal individual expressing the alternative strategy *x* = *z* + *δ* is given by

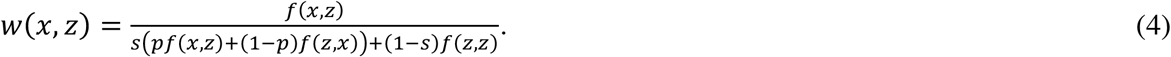

Note that the probability *p* is also the kin selection coefficient of relatedness *r* among individuals of species *A* within the focal patch (i.e. *r* = *p*; Bulmer 1994). The coefficient of relatedness gives the probability that an intraspecific social partner within a patch carries an identical allele relative to the population average.

We assume that the alternative helping strategy has a vanishingly small phenotypic effect, and therefore the selection gradient, denoted by *S*, is given by the slope of fitness on the phenotype (Otto and Day 2007). Thus, we have

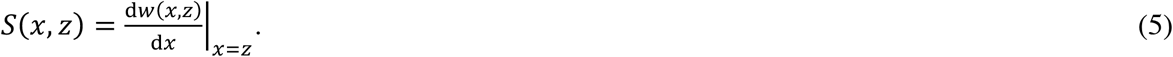

An evolutionarily stable helping strategy *z*^*^ is found when the selection gradient is null. That is, when there is neither selection for slightly more investment in helping nor selection for slightly less investment in helping, and therefore *S*(*z*^*^,*z*^*^) = 0 (Otto and Day 2007).

## Analysis and Results

Here we study how the different genetic and demographic factors affect the optimal investment into helping behaviour.

### Community lifespan

Let us focus on how the lifespan of the community, as given by *τ*, influences optimal investment in helping effort. Consistent with earlier work (Bull and Harcombe 2009), we find that mutualism is favoured at intermediate values of the community lifespan, while disfavoured when dispersal is early or late in the cycle of ecological development (Figure 2A). Interspecific cooperation slows down the initial growth rate of cooperators, and therefore, if dispersal occurs at this early stage (i.e. during the early exponential phase of bacterial growth), cooperation is not favoured by selection. On the other hand, interspecific competition is stronger at high densities when resources are scarce (i.e. during the stationary phase), and therefore cooperation is not favoured either when dispersal occurs at later stages of ecological development. This effect arises because the carrying capacity is fixed, and therefore as we approach the stationary phase the interaction between the two species becomes a zero-sum game. Any increase in the density of one species necessarily leads to a decrease in the density of the partner species. At intermediate community lifespans (i.e. during the exponential phase), interspecific cooperation is highest because sufficient time has elapsed for cooperators to recover from their initial slow growth and because interspecific competition is still weak. However, we find that competition imposed by intraspecific cheats (low *r*) is an important factor mediating this “window of opportunity” for interspecific cooperation, which can be drastically reduced when relatedness falls below 0.5 (Figure 2A).

**Figure 2.**
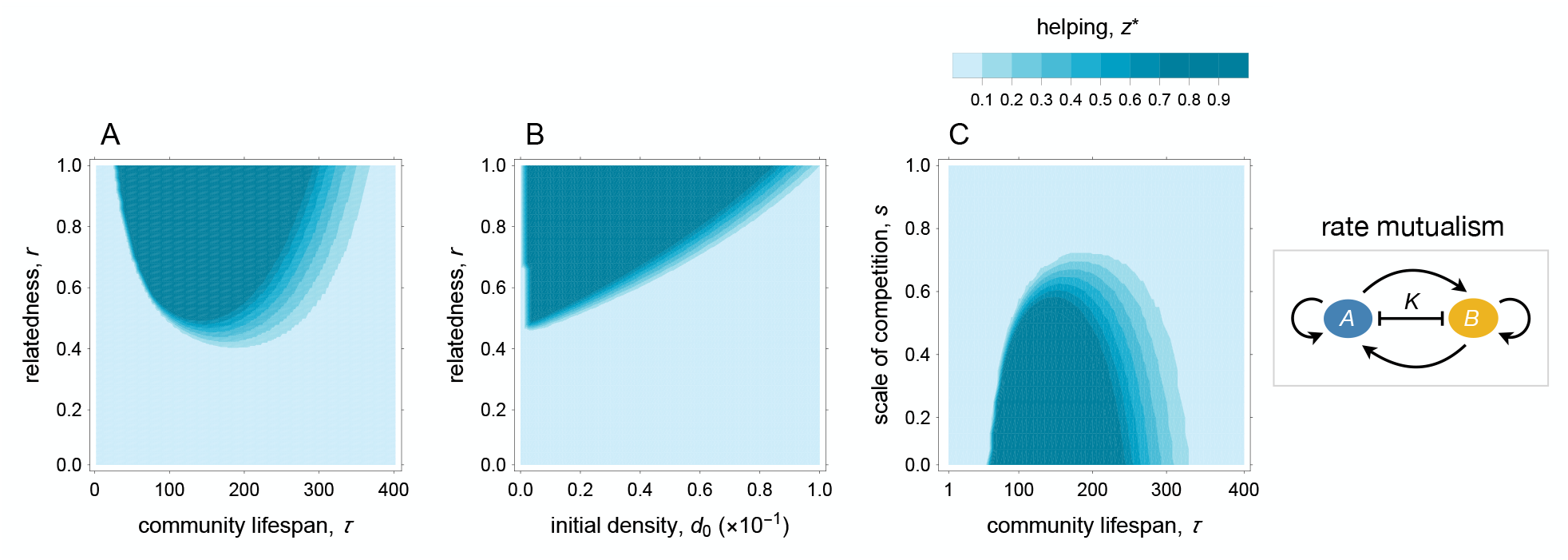
Rate mutualisms are favoured at high levels of relatedness and when dispersal occurs at intermediate values of community lifespan. (**A**) The optimal investment in helping (*z*^*^) as a function of community lifespan (*τ*) and relatedness (*r*). (**B**) The optimal investment in helping (*z*^*^) as a function of the initial density of bacteria (*d*_0_) and relatedness (*r*). (**C**) The optimal investment in helping (*z*^*^) as a function of the intensity of local competition (*s*) and community lifespan (*τ*). Parameter values: *ϕ*_A,0_ = 0.012, *ϕ*_B_ = 0.011, *α* = 0.01, ε= 10^−4^, *K* = 1, and (A) *s* = 0.5 and *d*_0_ = 0.01, (B) τ= 100 and *s* = 0.5, (C) *r* = 0.5 and *d*_0_ = 0.01.

### Abundance of colonisers

Let us now study the impact of the initial number *d*_0_ of colonisers on the evolution of cooperation. We find that cooperation is disfavoured as the initial number of colonisers increases (Figure 2B). As the initial density of colonisers increases, the amount of competition for local resources also increases. Therefore, even during the early growth stages, individuals are already suffering from intense competition for resources, which undermines the evolution of cooperation. As in the previous section, we find that intraspecific cheats have a strong impact on the evolution of interspecific cooperation, which can rapidly drop from full helping (i.e. *z*^*^ = 1) to no helping (i.e. *z*^*^ = 0) as soon as relatedness falls below 0.5.

### Scale of local competition

We now turn the attention to the impact of local competition on the evolution of cooperation. We find that as competition becomes more local (higher *s*), cooperation is less likely to evolve (Figure 2C). Moreover, under intense local competition, cooperation requires longer growth periods to evolve. Local competition, however, exerts less influence on the optimal cooperative strategies when the dispersal stage occurs at high densities (higher *τ*). This is because growth that occurs later in a community’s lifespan is strongly regulated by the availability of local resources (i.e. density-dependent local regulation), and therefore regulation of the population exerted through local competition becomes less important.

## Mutualism and Niche Expansion

So far, we have assumed a rate mutualism in which there is a growth rate cost for helpers, and a growth rate return benefit. This mirrors cases, for instance, of detoxification mutualisms where microbial species protect each other from antibiotics through the production of resistance enzymes that break down the antibiotics (e.g. beta-lactamases; Yurtsev et al. 2016). In many other cases, microbes secrete or excrete products that can lead to the expansion of the resource pool by providing access to novel substrates (see Figure 1B). Niche expansion can take various forms, including: the production of metabolic waste by one species that opens a new niche for a cross-feeding species (incidental cross-feeding; Hillesland and Stahl 2010, Estrela et al. 2012, LaSarre et al. 2017, Goldford et al. 2018); the secretion of exomolecules that enhance nutrient supply (e.g. extracellular enzymes and scavenging molecules; (e.g. D’Onofrio et al. 2010, Flint et al. 2012, Rakoff-Nahoum et al. 2016); or the killing of competitors that opens up space for growth and access to new resources (Yanni et al. 2019). Detoxification of environmental toxins can also lead to niche expansion, but it will often relieve growth inhibition as well. Here, we ask what is the role of niche expansion on the evolution of cooperation?

To study this scenario, we extend our base model to include a unidirectional cross-feeding relationship, impacting the yield of the community (see Figure 1B). Specifically, we now assume that the carrying capacity of the community is *K* = *K*_i_ + *K*_B_, where *K*_i_ is the intrinsic carrying capacity, and where *K*_B_ is the component of the carrying capacity that depends on the density of the partner species *B*. This is given by

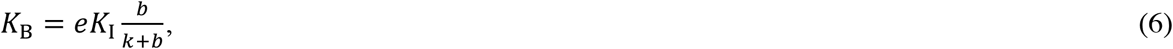

where *e* is a carrying capacity elasticity factor due to the presence of the partner species *B*, and *k* is a constant that determines how the carrying capacity increases with the density of the partner species (e.g. Frank 2010). If we set elasticity to zero (i.e. *e* = 0), we recover the base model, in which we have a rate mutualism with no niche expansion (i.e. *K* = *K*_I_, see Figure 3A and equations (1)-(3)).

**Figure 3.**
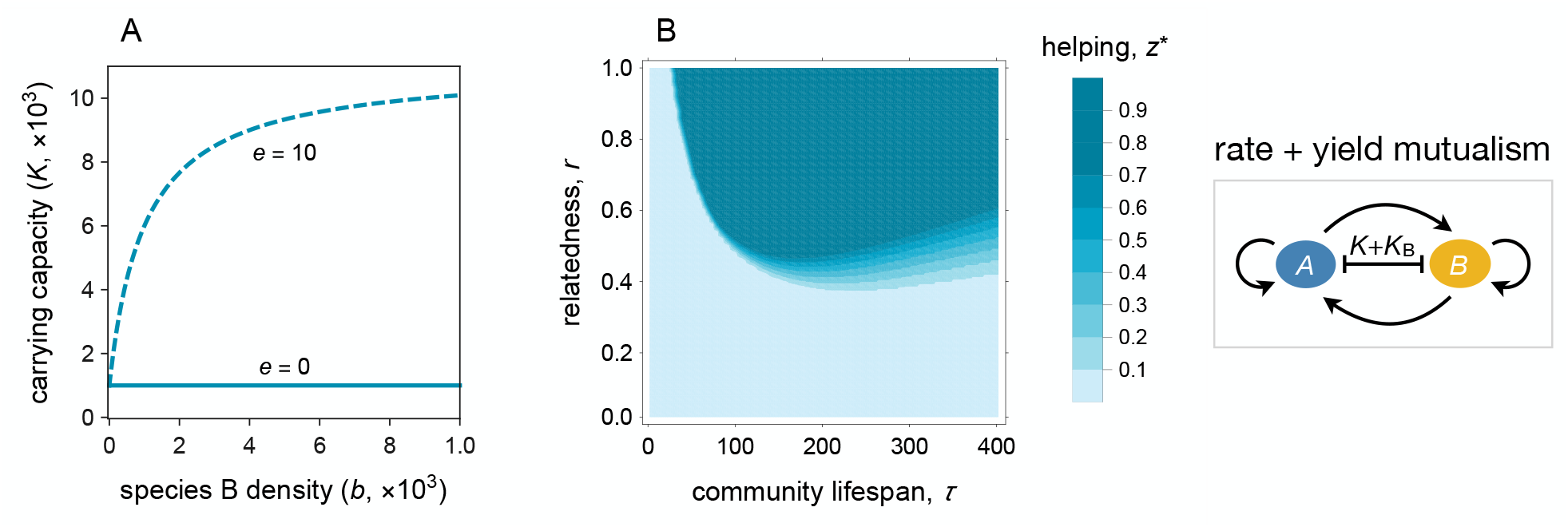
Niche expansion facilitates the evolution of mutualism at high densities. (**A**) Carrying capacity as a function of the partner species *B* density for various levels of elasticity (*e*). (**B**) Higher elasticity increases the window of opportunity within which cooperation is favoured by natural selection (see Figure 2A for the no elasticity baseline scenario), and may even favour cooperation at high densities. Parameter values: *ϕ*_A,0_ = 0.012, *ϕ*_B_ = 0.011, ε= 10^−4^, *s* = 0.5, *k* = 5, *α* = 0.01, *d*_0_ = 0.01, and *K*_i_ = 1.

In the absence of elasticity (i.e. *e* = 0), an increase in density of the partner species has a corresponding increase in the competition for resources to the focal cooperator and her relatives. Under this scenario, the additional competition for local resources generated by the partner species precludes the evolution of cooperation at high densities, as we have described above. If the elasticity *e* is sufficiently high, however, the partner species provides additional resources that expand the carrying capacity of the local patch. As a result, an increase in the density of the partner species need not lead to an increase in the competition for the local resources. Under this scenario, the additional resources provided by the partner species *B* offset the additional competition for resources, and therefore cooperation is favoured at high densities, as shown in Figure 3B (see also Figure 4, top row).

**Figure 4.**
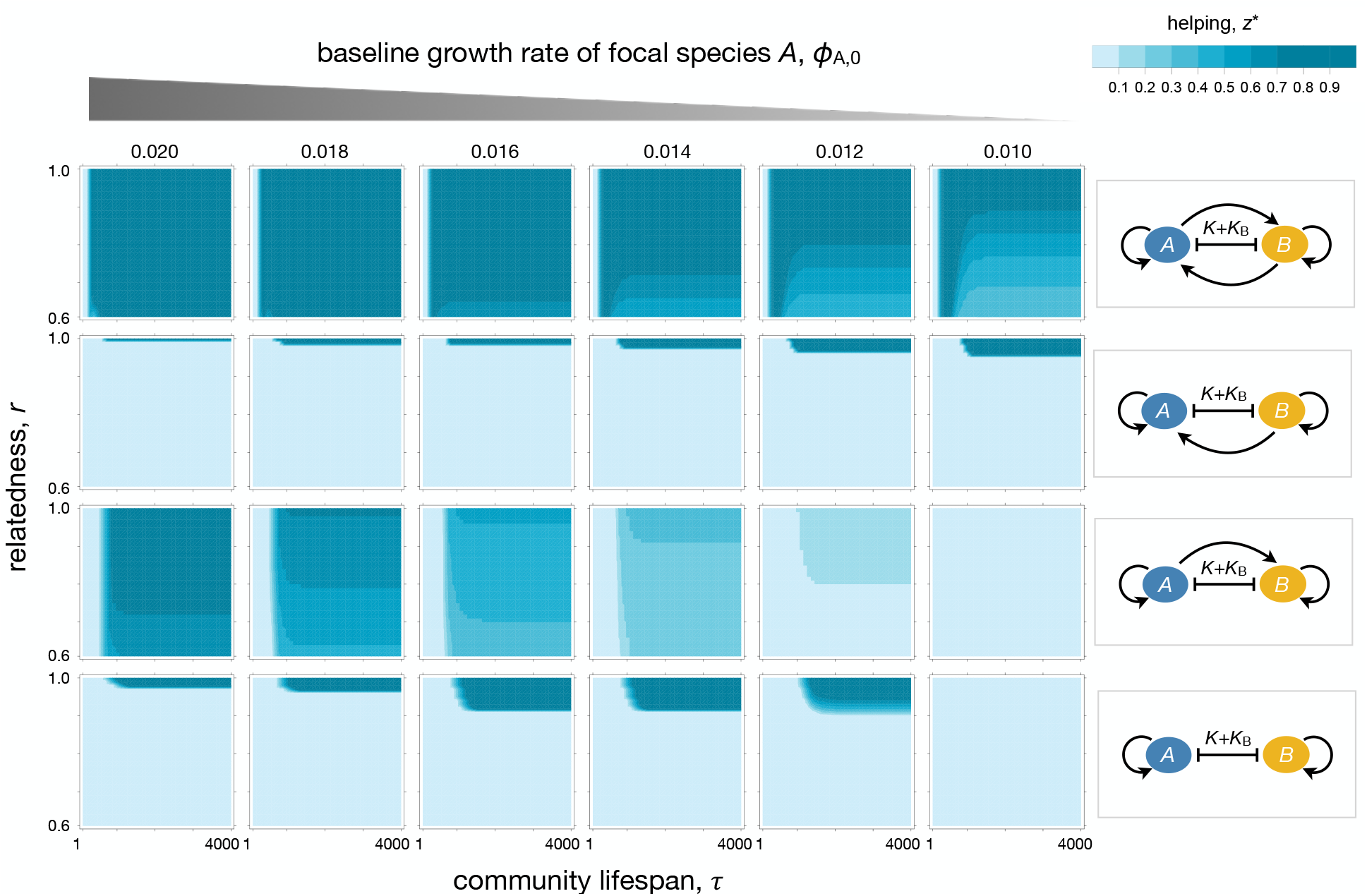
Rate mutualisms increase the range of parameters under which mutualisms can evolve. Optimal investment in helping (*z*^*^) as a function of community lifespan (*τ*) for various values of baseline growth rate (*ϕ*_A,0_) and rate of benefits (α and *β*_0_) in the presence of niche expansion. When there are cross-benefits (first row; *α* = 0.01, *β*_0_ = 0.01) mutualism evolves at intermediate and high densities. When there are return benefits (second row; *α* = 0.01, *β*_0_ = 0) mutualism evolves when communities are long-lived and is favoured by lower baseline growth rate of the focal species. When only the partner species enjoys a rate benefit (third row; *α* = 0, *β*_0_ = 0.01), interspecies cooperation becomes unstable, and only evolves when the baseline growth rate of the focal species A is sufficiently high. In the absence of rate benefits (fourth row; *α* = 0, *β*_0_ = 0), cooperation in the form of reproductive restraint evolves at high densities only, and is highest at intermediate value of baseline growth rate. Parameter values: *ϕ*_B_ = 0.011, ε= 10^−4^, *s* = 0.5, *k* = 5, *d*_0_ = 0.01, and *K*_i_ = 1.

So far, we have considered a scenario in which interspecific interactions influence the rate and the carrying capacity of each species simultaneously (i.e. *α* > 0, *β*_0_ > 0 and *e* > 0). As a result, we are unable to precisely gauge what are the forces driving the evolution of the mutualism. To investigate this, we now consider a simpler scenario in which interspecific interactions do not affect the growth rate of both the focal and the partner species (i.e. *α* = 0, *β*_0_ = 0), but the partner species still promotes niche expansion via the production of additional resources (*K*_B_). Under this scenario, we ask under what conditions natural selection favours the evolution of cooperation in the focal species. As shown in Figure 4 (bottom row), while cooperation does not evolve at intermediate densities, it does evolve at high densities. Unlike the previous scenario, here cooperation from the focal species *A* does not involve the secretion of any exo-products. Instead, cooperation takes the form of reproductive restraint in which individuals of the focal species *A* are selected to consume resources (replicate) more slowly (e.g. Kerr et al. 2006) to promote the growth of the partner species *B*. Slow growth has an initial cost for the focal species. However, this initial cost is offset by the niche expansion that occurs later on in the community lifecycle due to the additional individuals of the partner species *B*.

The evolution of reproductive restraint depends on the relative growth rate of the two interacting species (Figure 4). This growth-rate dependent outcome occurs because reproductive restraint has two contrasting effects on the fitness of the focal species. On the one hand, reproductive restraint has a fitness cost for the focal species because they are in direct competition with the partner species for limiting resources. On the other hand, reproductive restraint has a positive fitness effect for the focal species because it increases the abundance of the partner species, which also increases the carrying capacity of the local patch. When the baseline growth rate of the focal species is too small, the costs of reproductive restraint offset its benefits, and therefore, reproductive restraint does not evolve. If the baseline growth rate of the focal species is too high, the fast-growing focal species suppresses the growth rate of the partner species, which suppresses the benefits of cooperation at later ages of the community lifespan, and reproductive restraint is less favoured (Figure 4, bottom row).

We now consider a scenario in which in addition to the partner species leading to niche expansion, the focal species boosts the growth rate of the partner species (i.e. *α* = 0, *β*_0_ = 0.01). Do we still observe the evolution of interspecific cooperation? Under such scenario, we find that cooperation is less likely to evolve (Figure 4, third row). The cooperative behaviour of the focal species reduces its own growth (restraint), but it also increases the growth of the partner species. As a result, the additional growth rate of the partner species imposes an additional cost on the focal species. Overall, the benefits from niche expansion are not sufficient to offset the direct cost of cooperation arising from consuming common resources more slowly and the indirect cost of cooperation that arise from the additional competition imposed by the partner species.

Finally, let us now consider a scenario in which, in addition to niche expansion, the partner species also provides a growth rate benefit to the focal species (i.e. *α* = 0.01, *β*_0_ = 0). As shown in figure 4 (second row), we find that the focal species evolves reproductive restraint at high densities, but not at intermediate or low densities. Here restraint increases the density of the partner species, which in turn benefits the growth rate of the focal species, and also, the yield when at high densities owing to niche expansion by the partner species.

## Mutualism and Niche Overlap

So far, we have assumed that both species, *A* and *B*, compete for the same resources (represented by the common carrying capacity *K*). However, in many cases the degree to which niches of different species overlap varies. While in some cases species fiercely compete for common resources, i.e. complete niche overlap, in others they compete for some but not all resources, i.e. partial niche overlap, and yet in others they compete for very different resources, i.e. non-overlapping niches. We therefore extended our model to study the effect of niche overlap between species *A* and *B* on the evolution of mutualism. The degree to which species *A* and *B* compete for resources – degree of niche overlap – is modelled by an interspecific competition coefficient, denoted by *γ*, which varies from 1, i.e. complete niche overlap, to 0, i.e. non-overlapping niches. This coefficient affects the density-dependent terms in equations (1-3), such that they become: (1 – (*γ*(*a* + *a*_H_) + *b*) / *K*) in equation (1), which describes the growth trajectory of species *B*; and (1 – ((*a* + *a*_H_) + *γb*) / *K*), in equations (2) and (3), which describes the growth trajectories of species *A*. When there is complete niche overlap (*γ* = 1), we recover our initial model. However, when niches do not overlap (*γ* = 0), the density-dependent term of species *B* becomes 1 – *b*/*K*, and the density-dependent term of species *A* becomes 1 – (*a* + *a*_H_) / *K*.

Given niche expansion by the partner species (*K*_B_ > 0), relaxing competition for resources between the focal and partner species disfavours the evolution of reproductive restraint, and therefore yield-enhancing mutualisms are less likely to evolve (Figure 5 and Figure S2). Moreover, in such cases, mutualism with dispersal at later stages of community growth is no longer favoured when the degree of interspecific competition falls below 0.5 (Figure S2). This negative effect of lower interspecific competition on mutualism is especially pronounced in cases where the rate mutualism is turned-off (Figure S2, bottom row). Without niche expansion, cooperation at high densities is disfavoured regardless of the niche overlap (Figure 5). This suggests that interspecific cooperation in the form of reproductive restraint can only evolve when there is both a high degree of niche overlap and niche expansion.

**Figure 5.**
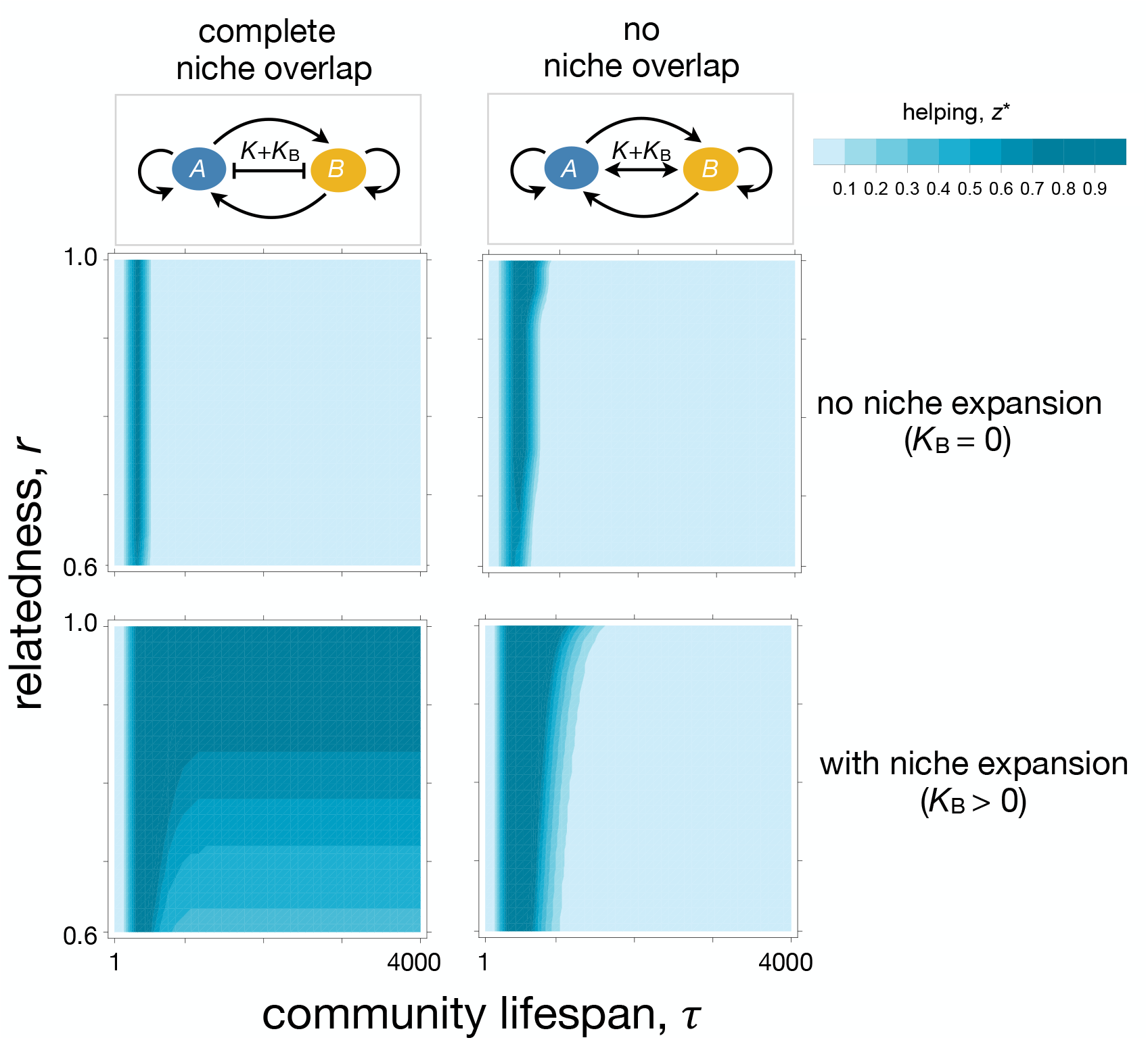
Non-overlapping niches disfavour the evolution of mutualism by reproductive restraint. Yield-enhancing mutualisms are sustained by reproductive restraint, which is only favoured under high degree of niche overlap. Rate-enhancing mutualisms, however, evolve regardless of the degree of niche overlap. Parameter values: *α* = 0.01, *β*_0_ = 0.01, *ϕ*_A,0_ = 0.014, *ϕ*_B_ = 0.011, ε = 10^−4^, *s* = 0.5, *k* = 5, *d*_0_ = 0.01, and *K*_i_ = 1.

## Discussion

In mutualistic interactions, individuals receive a benefit from their interspecific partners. In such cases, individuals should care for the survival and growth of their interspecific partners to the extent that this is associated with a higher probability of receiving the return benefit. In many cases, however, there is at least some degree of niche overlap between species, which may lead to interspecific competition for shared limiting resources, which, in turn, may have an adverse impact on the evolution of mutualisms. Previous theory seeking to understand this problem has shown that interspecific cooperation is more likely to evolve at intermediate densities (Bull and Harcombe 2009). This outcome arises because at intermediate densities the return benefits from interspecific cooperation are sufficiently high to outweigh the initial costs of cooperation and because the intensity of density-dependent competition for resources is still comparatively low.

The conclusion that mutualism evolves at intermediate densities relies on the assumption of a rate mutualism, in which individuals exchange resources that influence their growth rates. Mutualisms among microbes, by contrast, are commonly yield-enhancing (Samuel and Gordon 2006, Kolenbrander 2011, Ramsey et al. 2011). We have shown that in such cases interspecific cooperation can evolve even at high population densities, a condition that may be prevalent in natural microbial populations (e.g. Rakoff-Nahoum et al. 2016). In line with our findings, Rakoff-Nahoum et al. (2016) recently provided evidence of costly interspecific cooperation between Bacteroidales species that live at high densities in the human intestine. Specifically, *Bacteroides ovatus* secretes digestive enzymes that break down inulin, the products of which are then used by *Bacteroides vulgatus*, which, in turn, improves the fitness of *B. ovatus*. This study provides a rare example of a costly yield mutualism in a natural ecosystem. Evidence for costly mutualism between microbial species in a natural setting is still scarce, in part due to the difficulty of measuring the costs and benefits associated with a specific trait in complex environments such as the gut microbiota. Our study, however, suggests that rather than an exception, costly yield mutualisms may be a prevalent form of cooperative exchanges in the microbial world.

Understanding the nature of the costs and benefits accrued to individuals involved in interspecific exchanges is crucial for explaining the diversity and stability of mutualisms in microbial communities (e.g. Harcombe 2010, Rakoff-Nahoum et al. 2016, Harcombe et al. 2018). In rate mutualisms, the return benefits materialise quicker than in yield mutualisms, and so rate mutualisms are favoured at early stages of community growth (i.e. exponential phase of bacterial growth) while yield mutualisms are favoured at later growth stages (i.e. late-exponential and stationary phase). Given this difference in the rate of return of benefits, it is tempting to speculate that environmental conditions can predict the type of mutualism that is more likely to evolve in a given habitat. In general, we hypothesise that natural selection favours rate mutualisms in more variable and transient environments so that individuals maximise the exploitation of the available nutritional niches and minimise nutritional losses (e.g. due to resource washout). In contrast, in more stable and durable environments, natural selection should favour yield-enhancing mutualisms.

The evolution of interspecific cooperation also depends on the number of individuals who colonise a patch. If the density of colonisers is initially high, then there is intense competition for resources right from the beginning of colonisation, which narrows the window of opportunity within which cooperation evolves. Mixed genotypes in a patch, either due to mixed patch colonisation or due to mutations, may also disrupt cooperation, which may occur due to two processes. Firstly, collective action at an early stage of the demographic dynamics is required to produce enough benefits at intermediate stages, which can only happen if there is a sufficiently high number of cooperators during the early stages of community growth. Secondly, the benefits provided by the partner species at intermediate densities come in the form of public goods, and therefore all individuals of the focal species, both cooperators and cheats, benefit equally from the help provided by the partner species. As a result, high relatedness within the focal species ensures that the benefits of cooperation are directed towards cooperators, which prevents free-riders from exploiting the products of cooperation.

We find that the evolution of cooperation in the form of reproductive restraint in the focal species depends on the relative intrinsic growth rate of the two partner species. Reproductive restraint can evolve if the intrinsic growth rate of the focal species is high enough so that its own growth is not suppressed by the growth of its partner species at early stages of community development, but also not too high so that the focal species does not suppress the growth of its partner species, and consequently the benefits of cooperation that are accrued at later ages of the community lifespan. Broadly related to this idea, it has been shown that both growth rate differences between species and competition for a shared resource can affect the stability of cross-feeding interactions (Hammarlund et al. 2019). Starting with a bidirectional obligate cross-feeding mutualism between two species, Hammarlund et al. (2019) showed that when one partner becomes independent, coexistence can only be maintained when the obligate partner is the faster growing species but breaks down when the obligate partner is the slower grower. In our model, interspecific cooperation evolves between two species that can grow independently. It would be interesting to investigate whether restraint is more or less likely to evolve in systems where the partner species depends on the focal species for growth.

Our findings, so far, assumed that the focal and partner species compete for the same limiting resources, which is consistent with the idea of strong niche overlap. What happens if now they compete less, or not at all, for the same resources – that is, if there is low or no niche overlap? Previous work has suggested that low niche overlap and high relatedness can favour cooperation between species (Mitri and Foster 2013). Here we find that interspecific cooperation, in the form of reproductive restraint, is only favoured when interspecific competition is high, but not when interspecific competition is low. In other words, interspecific competition selects for interspecific cooperation. This effect occurs because without a shared limiting resource, the incentives for reproductive restraint are eliminated. Indeed, prudence in resource consumption no longer promotes the growth of the partner species and its associated yield-enhancing benefits, but it still carries the costs associated with intraspecific cheats. We would expect, however, low niche overlap to favour other forms of interspecific cooperation that directly impact the yield of partner species.

We have assumed that cooperation is constitutively expressed. Under this scenario, rate mutualisms are favoured at intermediate densities whilst yield mutualisms are favoured at high densities. These contrasting selective pressures raise the hypothesis that natural selection favours the density-dependent regulation of these cooperative traits. It is now well understood that intraspecific microbial cooperation is commonly regulated in a density-dependent manner (Xavier et al. 2011, Darch et al. 2012, Ghoul et al. 2016), often controlled by cell-cell signalling mechanisms termed quorum sensing (Whiteley et al. 2017). Quorum sensing limits the expression of multiple social traits to high-density environments, and limits the extent of intraspecific social cheating by positive-feedback control of cooperative behaviour (Allen et al. 2016). Our results suggest that interspecific cooperation may also be regulated in a manner that depends on total community density-for instance, through a mechanism akin to the interspecific quorum sensing molecule, the autoinducer AI2, that allows bacteria to determine their overall population density and regulate their behaviour accordingly (Xavier and Bassler 2005). Specifically, we predict that rate-enhancing traits are more likely to be up-regulated during earlier growth phases (low densities) but down-regulated during later growth phases (high density). By contrast, we predict that yield-enhancing traits are down-regulated during earlier growth phases but up-regulated during later growth phases and the stationary phase.

In this paper, we have not considered the co-evolutionary process between the two species and we have assumed that the partner species remains in its ancestral form and expresses a constant level of helping. Helping in the partner species is, however, also a trait under the action of natural selection, and allowing for the coevolution of helping between the two species may have important consequences for optimal levels of investment in cooperation (West et al. 2002). Moreover, we have also assumed that the partner species is not able to discriminate between cooperators and cheats in the focal species (Allen et al. 2016). For example, in the rhizobia-legume symbiosis, the plant is able to punish bacterial groups that do not help, and this mechanism has been shown to stabilise cooperation in this system (West et al. 2002). Also, here we assume that the focal and partner species are both autonomous – the most likely starting point of mutualisms. The production of costly leaky goods or services can, however, select for the evolution of one-way metabolic dependencies to save on metabolic costs (Morris et al. 2012) and ultimately interdependencies where microbes engage in the exchange of essential traits (Estrela et al. 2016). Extending our model in these multiple directions provides an interesting avenue for future studies.

## Supplementary Figures

**Figure S1.**
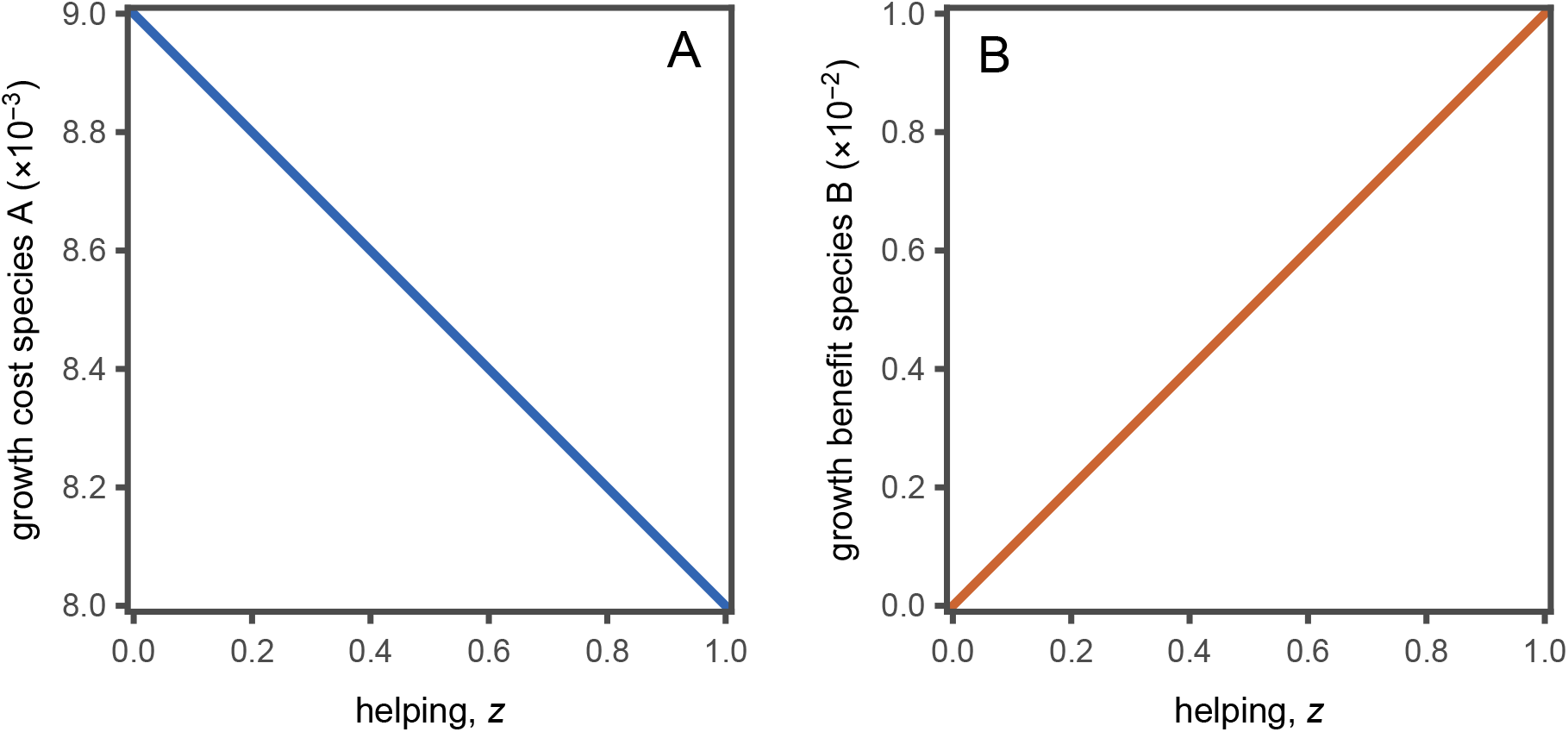
(**A**) Growth rate cost for the focal species as a function of their investment in helping (*z*). We assume *ϕ*_A_(*z*) = *ϕ*_A,0_ – *φ*_A_ *z.* (B) Growth rate benefit enjoyed by the partner species *B* as a function of the investment in helping (*z*) by species *A*. We assume *β*(*z*) = *β*_0_*z*. Parameter values: *ϕ*_A,0_ = 0.012, *φ*_A_ = 0.001, *β*_0_ = 0.01.

**Figure S2.**
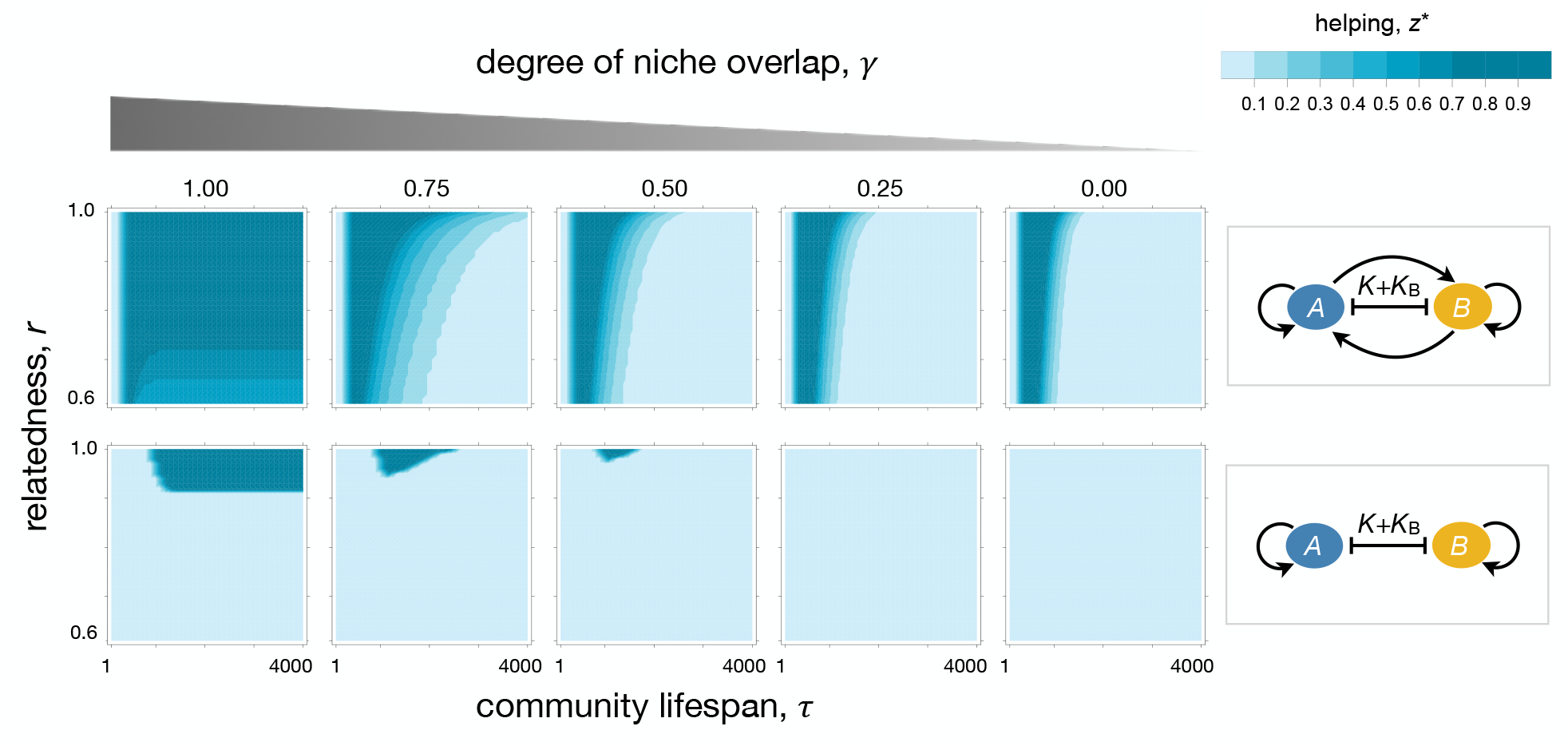
Low niche overlap disfavours the evolution of mutualism by reproductive restraint. When there are rate-enhancing cross-benefits (first row; *α* = 0.01, *β*_0_ = 0.01), little niche overlap (lower *γ*) selects against mutualism in long-lived communities but does not select against mutualism in short-and intermediate-lived communities. In the absence of rate benefits (second row; *α* = 0, *β*_0_ = 0), little niche overlap (lower *γ*) selects against mutualism in the form of reproductive restraint. Parameter values: *ϕ*_A,0_ = 0.014, *ϕ*_B_ = 0.011, ε = 10^−4^, *s* = 0.5, *k* = 5, *d*_0_ = 0.01, and *K*_i_ = 1.

